# Single-dose AAV-based vaccine induces a high level of neutralizing antibodies against SARS-CoV-2 in rhesus macaques

**DOI:** 10.1101/2021.05.19.444881

**Authors:** Dali Tong, Mei Zhang, Yunru Yang, Han Xia, Haiyang Tong, Huajun Zhang, Weihong Zeng, Muziying Liu, Yan Wu, Huan Ma, Xue Hu, Weiyong Liu, Yuan Cai, Yanfeng Yao, Yichuan Yao, Kunpeng Liu, Shifang Shan, Yajuan Li, Ge Gao, Weiwei Guo, Yun Peng, Shaohong Chen, Juhong Rao, Jiaxuan Zhao, Juan Min, Qingjun Zhu, Yanmin Zheng, Lianxin Liu, Chao Shan, Kai Zhong, Zilong Qiu, Tengchuan Jin, Sandra Chiu, Zhiming Yuan, Tian Xue

**Affiliations:** First Affiliated Hospital of USTC, School of Life Sciences, Division of Life Sciences and Medicine, University of Science and Technology of China, Hefei 230026, China; Neurodegenerative Disorder Research Center, CAS Key Laboratory of Brain Function and Disease, CAS Key Laboratory of Innate Immunity and Chronic Disease, Biomedical Sciences and Health Laboratory of Anhui Province, University of Science and Technology of China, Hefei 230026, China; State Key Laboratory of Virology, Wuhan Institute of Virology, Chinese Academy of Sciences, Wuhan 430071, China; Center for Biosafety Mega-Science, Wuhan Institute of Virology, Chinese Academy of Sciences, Wuhan 430071, China; Hefei Institutes of Physical Science, Chinese Academy of Sciences, Hefei 230031, China; Anhui Institute of Pediatric Research, Anhui Provincial Children’s Hospital, Hefei 230051, China; Chinese Academy of Sciences Center for Excellence in Brain Science and Intelligence Technology, Chinese Academy of Sciences, Shanghai 200031, China; Department of Clinical Laboratory, First Affiliated Hospital of Anhui Medical University, Hefei, 230022, China; Institute for Stem Cell and Regeneration, Chinese Academy of Sciences, Beijing 100101, China

**Keywords:** COVID-19 vaccine, AAV, SARS-CoV-2, variants, single dose

## Abstract

Coronavirus disease 2019 (COVID-19), which is triggered by severe acute respiratory syndrome coronavirus 2 (SARS-CoV-2) infection, continues to threaten global public health. Developing a vaccine that only requires single immunization but provides long-term protection for the prevention and control of COVID-19 is important. Here, we developed an adeno-associated virus (AAV)-based vaccine expressing a stable receptor-binding domain (SRBD) protein. The vaccine requires only a single shot but provides effective neutralizing antibodies (NAbs) over 598 days in rhesus macaques (*Macaca mulatta*). Importantly, our results showed that the NAbs were kept in high level and long lasting against authentic wild-type SARS-CoV-2, Beta, Delta and Omicron variants using plaque reduction neutralization test. Of note, although we detected pre-existing AAV2/9 antibodies before immunization, the vaccine still induced high and effective NAbs against COVID-19 in rhesus macaques. AAV-SRBD immune serum also efficiently inhibited the binding of ACE2 with RBD in the SARS-CoV-2 B.1.1.7 (Alpha), B.1.351 (Beta), P.1/P.2 (Gamma), B.1.617.2 (Delta), B.1.617.1/3(Kappa), and C.37 (Lambda) variants. Thus, these data suggest that the vaccine has great potential to prevent the spread of SARS-CoV-2.

---

Coronavirus disease 2019 (COVID-19) is a highly infectious respiratory disease that continues to pose a serious global public health emergency. The disease shows a high infection rate, long incubation period, and rapidly emerging variants, which have led to its rapid spread worldwide^1^. Many vaccines have been developed for the control of severe acute respiratory syndrome coronavirus 2 (SARS-CoV-2), the virus responsible for COVID-19, including vaccines based on messenger RNA (mRNA)^2, 3^, viral vectors^4, 5^, recombinant proteins^6^, and inactivated SARS-CoV-2^7, 8, 9^. Indeed, several of these vaccines have been shown to protect population from SARS-CoV-2 infection. However, most vaccines lack long-term protection efficacy^10^, and most of them require two or three injections to induce neutralizing antibodies (NAbs). Therefore, developing a vaccine that only requires single-dose immunization and provides long-term NAbs would be optimal for combating COVID-19.

Adeno-associated virus (AAV) is a single-stranded DNA parvovirus widely used for gene therapy and vaccines ^11, 12, 13^. AAV vector-mediated gene therapy products have been approved by the Food and Drug Administration (FDA) for the treatment of inherited blindness and spinal muscular atrophy^14^. AAV vectors have special features that are highly beneficial for clinical applications, such as low immunogenicity, long-lasting gene expression, safety, and high efficacy. Adenoviruses (AdVs) and AAVs are two types of viral vector used for gene delivery, with AdVs more commonly utilized for SARS-CoV-2 vaccines^5^. Both systems can infect a broad range of hosts, including dividing and non-dividing cells. However, there are several key distinctions between them, including onset and duration of gene expression, packaging capacity, and immune response. AdV vectors can accommodate larger inserts, but mediate transient protein expression and may cause severe inflammation and immune response. Compared with AdVs, AAVs exhibit longer lasting gene expression and lower immune response. Thus, we applied AAV vectors in the current study to develop a long-term expression vaccine for the prevention of COVID-19.

The SARS-CoV-2 spike protein mediates the binding of the virus to the human angiotensin converting enzyme 2 (ACE2) receptor for entry into target cells^15^. As such, it is the main antigen target for vaccines. Based on the SARS-CoV-2 spike protein structure (PDB: 6VXX), we found that the receptor-binding domain (RBD) was not as stable as the domain that spanned the spike protein from Q321 to S591, with the C and N tail forming a stabilizing beta-sheet (Figure 1a), hereafter termed SRBD. Thermal stability analysis also showed that the SRBD protein (56.88 ± 0.45 °C) was more thermostable than RBD (52.28 ± 0.77 °C) (Figure 1b). The AAV2/9 serotype was chosen as the vaccine carrier due to its high transduction efficiency in muscles. To assess the immunogenicity of the designed vaccines, we injected AAV-SRBD vaccines intramuscularly into both C57BL/6J and NIH mice at a dose of 1 × 10^11^ virus genomes (vg)/mouse, respectively (Figure S1a-b). AAV-CAG-GFP (1 × 10^11^ vg/mouse) was used as a control. Results showed that the AAV-SRBD could express well in the muscle of mice (Figure 1c). Moreover, AAV-SRBD resulted in high antibody titers in both NIH and C57BL/6J mice (Figure 1d and e). Thus, we used SRBD as the antigen for generating an AAV-based COVID-19 vaccine. To evaluate the tissue-specific expression patterns of AAV2/9, we analyzed the expression of AAV-CAG-GFP in several major mouse organs (Figure S1c-e). Green fluorescent protein (GFP) signaling was found in the injected muscle cells and liver cells of mice, but not in other major organs, i.e., heart, lung, spleen, kidney, and whole brain. Moreover, histological analysis illustrated that no significant pathological changes occurred in the major tissues of AAV-injected mice, e.g., lung, heart, liver, spleen, and kidney, compared with the naïve C57BL/6J mice (Figure S1f). These results suggest that the AAV-SRBD vaccine exhibited good immunogenicity and safety in mice.

**Figure 1.**
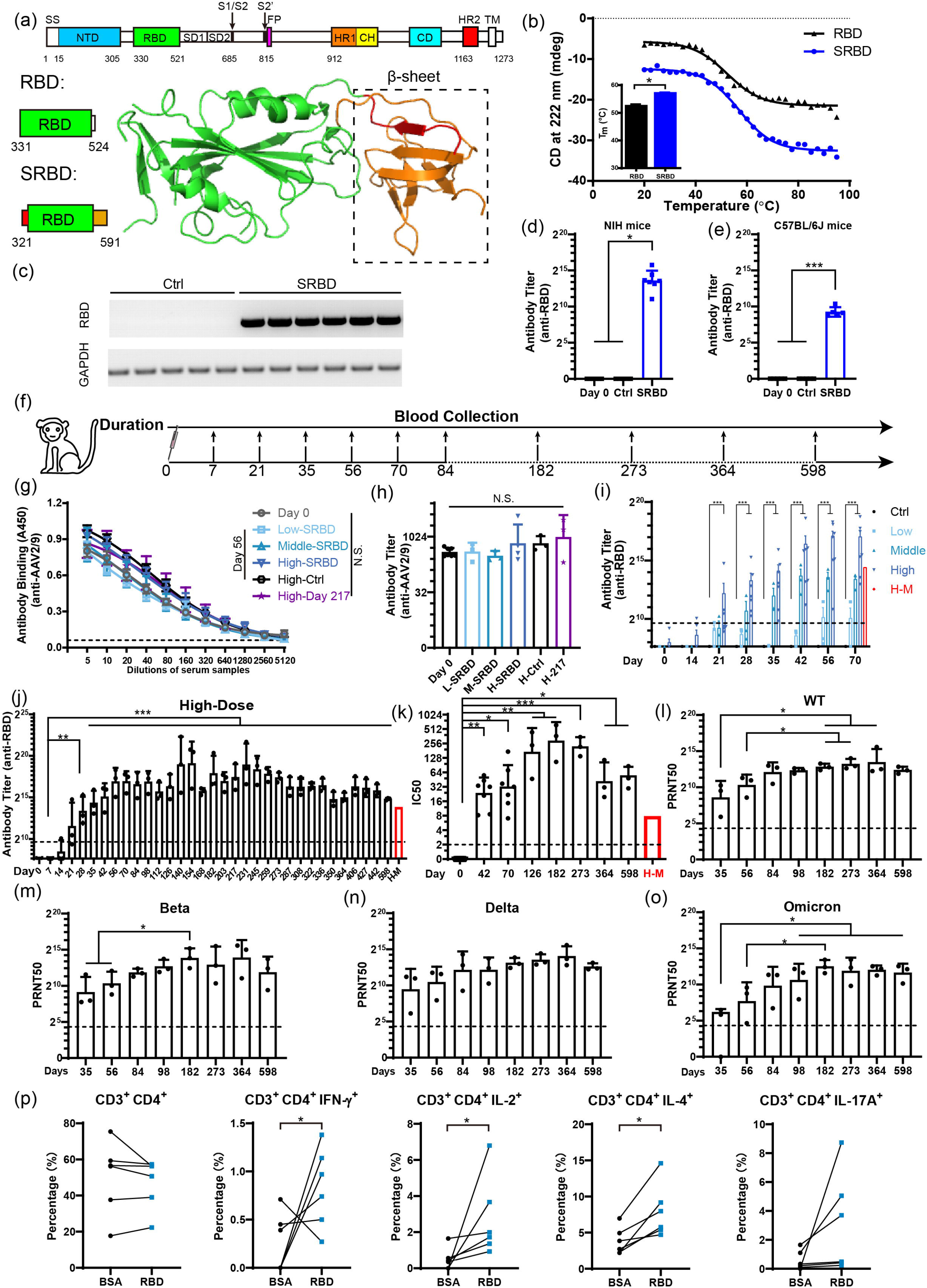
Single-dose SRBD vaccine elicits long-lasting and high-level humoral responses in rhesus macaques. (a) Schematic of RBD and SRBD structure. β-sheet was formed between red section (Q321 to P330) and orange section (C525 to S591), which provided stability to SRBD protein. SS, signaling sequence; NTD, N-terminal domain; RBD, receptor-binding domain; SD1, subdomain 1; SD2, subdomain 2; HR1, heptad repeat 1; CH, central helix; CD, connector domain; HR2, heptad repeat 2; TM, transmembrane domain. (b) Thermal stability analysis of RBD and SRBD proteins by circular dichroism spectroscopy. Thermal denaturation of RBD and SRBD are shown in black and blue curves, respectively, between 20 °C to 95 °C monitored at 222 nm. (c) mRNA expression of RBD in muscle samples of AAV-GFP-, AAV-SRBD-injected C57BL/6J mice at 28 dpv (n = 5 mice in control (Ctrl) group, n = 6 mice in SRBD group). (d) Quantitative analysis of RBD antibody titer calculated by ELISA at 28 dpv in NIH mice (n = 5 mice in Ctrl group, n = 7 mice in SRBD group, n=12 in Day 0 group). (e) Quantitative analysis of RBD antibody titer calculated by ELISA at 42 dpv in C57BL/6J mice (n = 5 mice in Ctrl and SRBD group, n=10 in Day 0 group). (f) Experimental strategy of AAV-SRBD vaccine analysis in rhesus macaques. (g) ELISA of AAV2/9 antibodies in macaque serum. (Day 0: serum collected before intramuscular injection; Low/Middle/High-SRBD: 56 dpv serum of low/middle/high-dose SRBD vaccine; High-Ctrl: 56 dpv serum of high-dose AAV-CAG-GFP control; High-Day 217: 217 dpv serum of high-dose SRBD vaccine; n = 3 macaques in Low-SRBD, Middle-SRBD, High-Ctrl, High-Day 217 groups; n = 7 macaques in Day 0 group; n = 4 macaques in High-SRBD group). (h) Quantitative analysis of AAV2/9 antibody titer of ELISA data in g. (i) Quantitative analysis of RBD antibodies titer calculated by ELISA from 0 to 70 dpv in low/middle/high-dose groups and AAV-CAG-GFP control macaques (n = 3 macaques in Ctrl, low-, and middle-dose group, respectively; n = 7 macaques in high-dose group). Mean RBD antibody titer of H-M is represented by red bar. The cutoff for the positive antibody titer is presented by black dot line. (j) Quantitative analysis of RBD antibody titer calculated by ELISA of RBD antibodies from 0 to 598 dpv in high-dose macaques of group 1(n = 3 macaques in each group). Mean RBD antibody titer of H-M is represented by red bar. The cutoff for the positive antibody titer presented by black dot line. (k) Quantitative analysis of RBD NAb IC50 levels calculated by competitive ELISA from 0 to 598 dpv in high-dose macaques (n = 7 macaques in day 0, 42 and 70 group; n = 3 macaques in other groups). Mean IC50 of H-M is represented by red bar. The cutoff for the positive IC50 presented by black dot line. (l) Serum PRNT50 level in group 1 high-dose macaques from 35 to 598 dpv calculated using plaque reduction neutralization test in Vero E6 cells against wild-type SARS-CoV-2 (n = 3 macaques in each group). The cutoff for the positive antibody PRNT50 presented by black dot line. (m) Serum PRNT50 level in group 1 high-dose macaques from 35 to 598 dpv calculated by plaque reduction neutralization test in Vero E6 cells against Beta variant (n = 3 macaques in each group). The cutoff for the positive antibody PRNT50 presented by black dot line. (n) Serum PRNT50 level in group 1 high-dose macaques from 35 to 598 dpv calculated by plaque reduction neutralization test in Vero E6 cells against Delta variant (n = 3 macaques in each group). The cutoff for the positive antibody PRNT50 presented by black dot line. (o) Serum PRNT50 level in group 1 high-dose macaques from 35 to 598 dpv calculated by plaque reduction neutralization test in Vero E6 cells against Omicron variant (n = 3 macaques in each group). The cutoff for the positive antibody PRNT50 presented by black dot line. Data (l to o) were obtained from one experiment. Each serum was tested three times as technical replicates (p) Percentage of CD3^+^CD4^+^, CD3^+^CD4^+^IFN-γ^+^, CD3^+^CD4^+^ IL-2^+^, CD3^+^CD4^+^ IL-4^+^, and CD3^+^CD4^+^ IL-17A^+^ cells in blood of high-dose rhesus macaques activated by BSA or RBD peptide (n = 6 macaques in each group). Values are means ± SEM or geometric mean + geometric standard deviation for antibody titer and PRNT50. *: *P* < 0.05; **: *P* < 0.01; ***: *P* < 0.001.

To further examine vaccine safety and efficacy, we tested the AAV-SRBD vaccine in a nonhuman primate (NHP) species. Two groups of rhesus macaques (*Macaca mulatta*) were used for the study. The first group included three macaques with high-dose vaccine (1×10^12^ vg/macaque) and was used for long-term monitoring of the NAbs titers. The second group included 13 macaques with different doses. 13 macaques were randomly divided into three groups, then received a single-dose immunization of 1×10^12^ vg/macaque (high-dose, four macaques), 1×10^11^ vg/macaque (middle-dose, three macaques), or 1×10^10^ vg/macaque (low-dose, three macaques) of AAV-SRBD, respectively. AAV-CAG-GFP (1×10^12^ vg/macaque) was used as the control (three macaques) (Table S1). The dosage dependent effect of the vaccine, the body weight, antibody titer, pathological indicators in blood and hepatic function of macaques in the second group were examined until 70 post vaccination (dpv). All intramuscular-injected macaques in the second group showed normal body weight post injection (Figure S2a). Blood samples from all macaques were collected to assess the antibody titers (Figure 1f). One potential limitation of AAV vaccine application is that most humans and macaques have experienced wild-type AAV exposure, which can result in pre-existing AAV antibodies and inhibition of AAV transduction in primates^16^. Given this, we estimated the levels of AAV2/9 antibodies in the macaques before and after vaccination. First, we randomly selected seven macaques, including two macaques each in the low- and middle-dose groups and three macaques in the high-dose group (group 1) to evaluate their pre-existing levels of AAV2/9 antibodies. All tested macaques were AAV2/9 antibody-positive (Figure 1g and h), suggesting that AAV2/9 antibodies may commonly exist in this species. Interestingly, the AAV2/9 antibodies levels did not change significantly from 56 to 217 dpv compared with day 0, even in the high-dose AAV-SRBD macaques. These results suggest the pre-existence of AAV2/9 antibodies in macaques before immunization, and that intramuscular injection of the AAV-based vaccine did not boost AAV antibody levels.

Even though AAV2/9 antibodies pre-existed in the macaques, the seroconversion rate (antibody titer > 800) reached 100% on 35 dpv in the high- (7/7) and middle-dose (3/3) macaques, but only 33.3% (1/3) at 56 dpv in the low-dose macaques (Figure 1i). Accordingly, the AAV-SRBD vaccine demonstrated good immunogenicity in the high- and middle-dose macaques, but not in the low-dose macaques, as tested by enzyme-linked immunosorbent assay (ELISA) (Figure 1i and Figure S2c) and competitive ELISA (Figure S2d-g). The SRBD NAbs in the high- and middle-dose macaques effectively inhibited interactions between the RBD and ACE2, and efficacy was better than that of mixed sera from convalescent COVID-19 patients with severe disease (H-M) (Figure S2d-g). These results suggest that AAV-SRBD induced robust humoral responses in NHPs, and vaccine efficacy appeared to be highly dose-dependent (*P* = 0.0293 in Figure 1i; *P* = 0.0221 in Figure S2e; *P* = 0.0090 in Figure S2g by one-way ANOVA). To assess the long-term humoral immune response of the AAV vaccine, we also monitored the SRBD antibody levels in the high-dose macaques (group 1) from days 0 to 598 dpv (Figure 1j). Results indicated that SRBD antibodies emerged on 21 dpv in the high-dose macaques and remained at a high level until 598 dpv, with an average titer higher than found in the H-M^17, 18^. The absorbance at 450nm (A450) values demonstrated that the binding of RBD to SRBD antibodies increased with time but decreased slightly at 364 dpv and 598 dpv (Figure 1j and S2h), as found for the inhibitory ability of NAbs (Figure 1k and S2i). However, the inhibition rate was still higher than that in the H-M samples at 598 dpv. These results indicate that AAV-SRBD triggers a robust and long-lasting humoral response after a single-dose of vaccine, and that pre-existing AAV2/9 antibodies do not interfere with the vaccine immunity.

The SRBD NAbs from high dose macaques (group 1) were further measured using the plaque reduction neutralization test (PRNT), which is the gold-standard for determining immune protection. As the SARS-CoV-2 variants especially Delta and Omicron variants widely spread worldwide, we evaluated the SRBD NAbs against authentic wild-type SARS-CoV-2, Beta, Delta and Omicron variants using standard PRNT assay (Figure 1l-o, Figure S3). The 50% reduction in plaque count (PRNT50) value of the SRBD NAbs increased from 35 to 182 dpv, and remained at a high level (geometric mean of PRNT50 >1:2 048) in all macaques after 98 dpv against the wild-type SARS-CoV-2 virus. The PRNT50 values against the Beta and Delta variants were similar to the value against the wild-type SARS-CoV-2 virus (geometric mean of PRNT50>1:2 048) after 98 dpv. However, geometric mean of PRNT50 against the Omicron variant was lower than that against wild-type SARS-CoV-2 virus, but all sera tested still kept in high level, showing a PRNT50 from 893 to 11 112 after 182 dpv. These data together illustrated that the AAV-SRBD vaccine induced high and effective NAbs against the wild-type SARS-CoV-2, Beta, Delta and Omicron variants in rhesus macaques, and the AAV-SRBD vaccine provided efficient cross neutralization against major SARS-CoV-2 variants. Based on these results, it is reasonable to believe that our vaccine is broad spectrum and could provide protection for future emerging variants.

To assess the antigen-specific T cell responses to the AAV-SRBD vaccine, we used the RBD peptide pool to stimulate peripheral blood mononuclear cells (PBMCs) collected from high-dose macaques at 35 dpv. Compared to the bovine serum albumin (BSA) control, the percentages of the CD4^+^ IFN-γ^+^, CD4^+^ IL-2^+^, CD4^+^ IL-4^+^, and CD8^+^IL-4^+^ T cells increased under RBD peptide pool stimulation (Figure 1p and Figure S4). These results indicate that RBD-specific Th1 (IFN-γ^+^ and IL-2^+^) cell and Th2 (IL-4^+^) cell responses in PBMCs can be activated by stimulation of the RBD peptide pool after vaccination.

The toxicity of the SRBD vaccine was further evaluated in rhesus macaques. As of 598 dpv, no deaths, impending deaths, or significant abnormalities in clinical physiology were found in any macaque in group 1. Widely analyzed pathological indicators also showed that lymphocyte subgroup (CD20^+^, CD3^+^, CD3^+^CD4^+^, and CD3^+^CD8^+^) distribution was normal before and after intramuscular injection (Figure S5, stimulation with cocktail) by 70 dpv. These results strongly suggest that the AAV-SRBD vaccine is safe and does not trigger severe inflammation. Other pathological indicators in blood, i.e., white blood cell (WBC), monocyte (MONO), neutrophil (NEUT), eosinophil (EO), basophil (BASO), lymphocyte (LYMPH), red blood cell (RBC), and platelet (PLT) counts, were also normal before and after immunization with different doses of the AAV vaccine (Figure S6). These results suggest that the vaccine did not cause blood toxicity after injection. On the other hand, we found that the AAV was expressed in the liver of C57BL/6J mice (Figure S1c), which was also supported by other studies^19, 20, 21^. Similarly, we detected SRBD expression in the livers of the high-dose macaques, but very little in the low-dose group. To exclude the hepatotoxicity of the AAV-SRBD vaccine, we examined the hepatic function of the immunized macaques. Results showed no significant change of the levels of Alkaline phosphatase (ALP), Total bilirubin (TBIL), Alanine aminotransferase (ALT), Aspartate aminotransferase (AST) before and after vaccination. Thus, although AAV gene expression was detected in the liver of the high-dose macaques, it did not appear to trigger liver dysfunction and inflammation (Figure S7).

Currently, several SARS-CoV-2 variants are of concern worldwide. Thus, we investigated whether the SRBD vaccine shows efficacy against variants such as B.1.1.7 (Alpha), B.1.351 (Beta), P.1/P.2 (Gamma), B.1.617.2 (Delta), B.1.617.1/3 (Kappa), and C.37 (Lambda). To test the inhibitory ability of SRBD NAbs against different SARS-CoV-2 variants, we generated several RBDs in different variants, i.e., B.1.1.7, B.1.351 and P.1/P.2, B.1.617.1/2/3, and C.37. Based on competitive ELISA, the macaque serum effectively inhibited interactions between the RBD mutants and ACE2 (Figure S8). Therefore, the SRBD NAbs appear to offer long-term inhibitory activity against SARS-CoV-2 variants. These results indicate that the SRBD vaccine has the potential to block infection from SARS-CoV-2 variants.

In conclusion, we developed a single-dose vaccine that can provide long-term protection against SARS-CoV-2. Our results showed that SRBD is more thermostable than RBD. The AAV-SRBD vaccine could induce good seroconversion rate in both NIH and C57BL/6J mice. This vaccine overcomes the multiple injection requirement of current vaccines and provides high-level and long-lasting RBD NAbs. The presence of pre-existing immunity to AAV2/9 here did not restrict the delivery or efficacy of AAV-SRBD. We suspect that AAV rapidly enters the cells and AAV antibody titer is relatively low in muscles, allowing AAV-SRBD to overcome the inhibition of AAV antibodies. A potent immune response to AAV-ovalbumin was observed when AAV was administered intravenously but not when administered intramuscularly.^22^ Importantly, the SRBD vaccine provides cross neutralization against emerging variants. Further studies on wild type SARS-CoV-2 and Omicron variant challenge in macaques are in progress to explore the protective immunity of the AAV-SRBD vaccine. Another study reported on two AAV-based vaccines that demonstrate long NAb durability in mice and NHPs with a single injection, further supporting the safety and efficacy of AAV-based vaccines^23^. Thus, this vaccine shows great potential for the control of COVID-19.

## Materials and Methods

### Animal husbandry samples

C57BL/6J mice, NIH mice, and rhesus macaques (*Macaca mulatta*) were maintained at 25 °C on a 12 h:12 h light:dark cycle in an animal room at the University of Science and Technology of China. All procedures were conducted in accordance with the Principles for the Ethical Treatment of Animals approved by the Animal Care and Use Committee at the University of Science and Technology of China (Animal ethics number: 202006220919000464981). The macaques received a single-dose immunization (1 mL) of SRBD (1 × 10^12^ vg/macaque (high-dose, 4 males and 3 females), 1 × 10^11^ vg/macaque (middle-dose, 2 males and 1 female), or 1 × 10^10^ vg/macaque (low-dose, 1 male and 2 females) or AAV-CAG-GFP (1 × 10^12^ vg/macaque; control, 1 male and 2 females) by intramuscular injection. Mice received a single-dose immunization (20 μL) of SRBD, RBD, or AAV-CAG-GFP (1 × 10^11^ vg/mouse, males and females). For immunohistochemical analysis, mice received different doses (1 × 10^11^, 1 × 10^10^, and 1 × 10^9^ vg/mouse, males and females) of AAV-CAG-GFP.

### AAV-package

The AAV vaccines were packaged in HEK-293T cells. In brief, 10 μg of pHelper vector, 5 μg of AAV2/9 vector, and 5 μg of pITR vector (RBD/SRBD/GFP, Figure S1a) were transfected into a 10-cm diameter dish with HEK-293T cells by polyethylenimine (PEI) (PolyScience, Niles, USA). The HEK-293T cells were cultured with Dulbecco’s Modified Eagle Medium (DMEM, Gibco, 11965-092) with 10% fetal bovine serum (FBS) (Gibco, 16000-044) at 37 °C. The supernatant of the HEK-293T cells was harvested at days 3 and 6 after transfection. The supernatant was concentrated with Ultra-15 Centrifugal Filters (Millipore, UFC905024) and then gradient-purified with 15% to 60% Optiprep (Sigma, D1556). The AAV vaccine was harvested and washed using phosphate-buffered saline (PBS) six times, then diluted in 150 μL of PBS. The titers of the AAV vaccines were calculated by quantitative real-time polymerase chain reaction (qRT-PCR). The AAV vaccines were stored at −80 °C.

### Electron microscope (EM) sample preparation

The AAV vaccines (1 × 10^12^ vg/mL) were added to carbon-coated copper grids previously glow-discharged at low air pressure and stained with 2% uranyl acetate for 90 s. The EM was operated at an acceleration voltage of 120 kV. Images were recorded using a Tecnai G2 Spirit 120kV EM at 23 000× magnification.

### Protein expression and purification

The methods for purifying the SARS-CoV-2 RBD [amino acid (AA) 321–591], SARS-CoV-2 RBD variants, and human ACE2 extracellular domain (AA 19 to 615) followed previous research ^18^. In brief, target genes were inserted into the pTT5 vector, which contains a IFNA1 signal peptide at the N-terminus and a TEV enzyme site connected to the human IgG1 Fc at the C-terminus. The expression vectors were then transiently transfected into HEK-293F cells using polyethyleneimine (Polyscience). After 3 days, the supernatant was collected by centrifugation at 5 000 ×*g* for 15 min at 4 °C. About ¼ volume of 1 × PBS was added to adjust the pH of the supernatant. The supernatant was then loaded onto the protein A column and the target protein was eluted with 0.1 M acetic acid on ÄKTA pure (GE Healthcare). The collected protein was added to 1 mM edetate disodium (EDTA), 5 mM dithiothreitol (DTT), and tobacco etch virus (TEV) protease to remove Fc on a shaker in a 4 °C freezer. After dialysis in 1 × PBS, tandem protein A and nickel columns were used for further purification. Both Fc and undigested protein were loaded onto the protein A column and TEV (6 × His tag) was loaded onto the nickel column. The target protein was collected during flow through.

### Thermal stability analysis

To compare the thermal stability of RBD and SRBD, circular dichroism (CD) spectra were acquired on a Chirascan Spectrometer (Applied Photophysics, Leatherhead, UK). Prior to CD measurements, the sample buffers were changed to PBS and the protein concentration was adjusted to 0.5 mg/mL, as determined by its absorbance at 280 nm. For thermal titration, CD spectra were acquired from 20 to 95 °C with temperature steps of 5 °C and wavelengths between 180–260 nm. The CD signals at 222 nm were used to characterize structural changes during thermal titration. The data were fitted by Prism v6 to calculate the Tm values.

### Vaccine immunogenicity analysis

The C57BL/6J and NIH mice (8 weeks old; male and female; 20–25 g body weight for C57 mice and 30–35 g body weight for NIH mice) were randomly divided into five groups (five mice per group). The mice were intramuscularly injected with RBD or SRBD AAV vaccines or the AAV-CAG-GFP control at a dose of 1 × 10^11^ vg (20 μL). The macaques were randomly divided into four groups (seven macaques in high-dose group and three macaques in other groups) and intramuscularly injected with 1 mL of SRBD vaccine (1 × 10^12^, 1 × 10^11^, and 1 × 10^10^ vg/macaque for high/middle/low dose, respectively) or AAV-CAG-GFP control (1 × 10^12^ vg/macaque). Blood was collected from macaques before immunization and at every 7 days before day 42 after injection and every 14 or 21 days after day 42.

### ELISA and competitive ELISA

Nunc MaxiSorp plates were coated with 3 μg/mL recombinant RBD, P.1/P.2 RBD, B.1.1.7 RBD, B.1.617RBD, B.1.351 RBD, or 3 μg/mL AAV9 at 4 °C overnight. After washing four times with PBS (3 min each time), the plates were blocked with 5% non-fat milk in PBS at room temperature for 2 h. Serially diluted serum (5% non-fat milk in PBST (PBS with 0.1% Tween-20) for ELISA or 5% non-fat milk in PBST with 15 nM biotin-ACE2-TEV-Fc for competitive ELISA) was added to the plates, which were then incubated at room temperature for 1 h. After washing three times with PBST (3 min each time), horseradish peroxidase (HRP)-conjugated goat anti-mouse IgG (Sangon Biotech, D110087, mouse serum, ELISA), rabbit anti-monkey IgG (Cellwaylab, C020217, macaque serum, ELISA), or HRP-conjugated streptavidin (Beyotime, A0303, competitive ELISA) were added, followed by incubation at room temperature for 1 h. For macaque serum ELISA, HRP-conjugated goat anti-rabbit IgG (Sangon Biotech, D110058) was added, followed by incubation at room temperature for 1 h. After washing three times with PBST (3 min each time), TMB substrate (Beyotime, P0209) was added for 8 min, then stopped by 1 M H_2_SO_4_. Absorbance at 450 nm was measured with a microplate reader.

Antibody titer was calculated as the dilution of the serum that induces an A450 value twice that of the A450 value of the negative control. Dotted lines in Figure 1g and Figure S2b and h represent the A450 value of the negative control. The Dotted line in the Figure 1i-l and Figure S2f represents the cutoff of RBD antibody titer and IC50 of the macaques’ sera (Cutoff defined as 4-fold higher in antibody titer or IC50 compared to the negative baseline; because of the antibody titer of negative baseline could not been calculated by the Elisa curve as it is much lower than the first dilution of serua, so we use the the first dilution (1:200 in Elisa and 1:1 in competitive ELISA) as the antibody titer of negative baseline).

### Virus and cells

Vero E6 cells were maintained in Dulbecco’s modified Eagle’s medium (DMEM, Gibco) supplemented with 10% fetal bovine serum (FBS, ExCell Bio) and 1% penicillin-streptomycin (Gibco) at 37°C under a 5% CO2 atmosphere. The SARS-CoV-2 WIV04 strain was originally isolated from a COVID-19 patient in 2019 (GISAID, accession no. EPI_ISL_402124)^24^; Beta variant (NPRC2.062100001) was kindly provided by Chinese Center for Disease Control and Prevention^25^, and Delta variant (B.1.617.2; GWHBEBW01000000) by Prof. Hongping Wei; Omicron variant (B.1.1.529; BA.1) was isolated from a throat swab of a patient from Hong Kong by the Institute of Laboratory Animal Sciences, Chinese Academy of Medical Sciences (CCPM-B-V-049-2112-18). All processes in this study involving authentic SARS-CoV-2 were performed in a BSL-3 facility.

### Plaque reduction neutralization test in Vero E6 cells

The neutralizing titer was determined as describe previously with slight modification^26^. Briefly, antibodies were serially diluted with DMEM containing 2.5% FBS, and mixed with equal volume of virus suspension and incubated at 37°C for 1 h. The mixture was added to Vero E6 monolayer cells in 24-well plates and incubated for another 1 h, and the inoculate was replaced with DMEM containing 2.5% FBS and 0.9% carboxymethyl-cellulose. The plates were fixed with 8% paraformaldehyde and stained with 0.5% crystal violet 3 days later. Plaque reduction neutralizing titer was calculated using the “inhibitor vs normalized response (Variable slope)” model in the GraphPad Prism 8.0 software. The Cut-off value is calculated by the negative control (geometric mean + 3 times of geometric standard deviation).

### RT-PCR

Total RNA was extracted from organs with Trizol reagent (Invitrogen, 15596026) and a PrimeScript RT Reagent Kit (Takara, RR037A). Forward and reverse primers were designed to target RBD sequence (forward: 5’- GTGTACGCCTGGAATCGGAA -3’ and reverse: 5’- GATCTCGGTGCTGATGTCCC -3’).

### Hematoxylin-eosin (H&E) staining

Mice were anesthetized with sodium pentobarbital (40 mg/kg, intraperitoneal injection), then perfused with PBS and fixed in 4% paraformaldehyde (PFA). Organs were post-fixed in 4% PFA overnight at 4 °C, then dehydrated in 15% and 30% sucrose, respectively. Organs were sectioned at a thickness of 10 μm for H&E staining with a freezing microtome. Isopropanol (500 μL) was added to the slices and incubated at room temperature for 1 min. The slices were air dried, stained with hematoxylin (Agilent, S330930-2, 1 mL), and incubated at room temperature for 7 min. The slices were then washed 10 times (10 s each time) with ultrapure water, followed by the addition of bluing buffer (Agilent, CS70230-2, 1 mL) and incubation at room temperature for 2 min. After washing five times with ultrapure water (10 s each time), eosin mix (Sigma, HT110216, 1 mL) was added, and the sections were incubated at room temperature for 1 min.

### Immunohistochemical analysis

Organs (muscle, liver, heart, lung, spleen, kidney and brain) were sectioned (40-μm thick) for immunohistochemical analysis with a freezing microtome. After washing with PBS three times (5 min each time), the slices were blocked with 3% bovine serum albumin (BSA) and 0.1% Triton-X100 in PBS for 1 h at room temperature. The slices were then stained using 1:1 000 anti-GFP antibody (Earthox, E002030-02) in blocking buffer overnight at 4 °C. Slices were washed with PBS three times (15 min each time) and incubated with secondary antibodies Alexa Fluor 488 donkey anti-mouse IgG (1: 1000, Thermo Scientific, A21202) for 2 h at room temperature. DAPI (1:1000, Thermo Scientific, D3571) was used to stain cell nuclei. Confocal images were captured using a Leica microscope.

### Peripheral blood mononuclear cells (PBMCs), exocellular and intracellular staining, and flow cytometry

Blood samples from macaques were collected in EDTA-2K tubes. Ficoll medium (3 mL, Solarbio, P4350) was first added into a 15-mL tube, followed by blood (3 mL) and density gradient centrifugation at 400 ×*g* for 20 min at room temperature (ACC/DEC: 6/2). The plasma was then collected and stored at −80 °C. CELLSAVING buffer (Xinsaimei, C40100) was used to resuspend the PBMCs after thorough washing with PBS, with the cells then stored at −80 °C.

The frozen cells were resuspended and washed in RPMI medium 1640 (Gibco). Anti-CD3 (BD Biosciences, 557705), anti-CD4 (BioLegend, 357423), anti-CD8 (BioLegend, 301007), and anti-CD20 (BioLegend, 302310) antibodies were added to the cells for staining for 30 min in the dark on ice. After washing with PBS (30 s), the cells were tested on a BD FACSVerse flow cytometer. The PBMCs were also resuspended in RPMI medium 1640, with a cocktail (BD Biosciences, 550583) added to activate the cells at 37 °C for 4 h. Cells were washed with 1 × PBS (30 s) and stained on ice in the dark for 30 min with anti-CD3 (BD Biosciences, 557705), anti-CD4 (BioLegend, 357423), and anti-CD8 (BioLegend, 344714). The cells were then fixed and permeabilized using a Cytofix/Cytoperm Soln Kit (BD Biosciences, 554714). Afterwards, cells were stained with anti-IFN-γ (BioLegend, 502526), anti-TNF-α (BioLegend, 502930), and anti-IL-10 (BioLegend, 501420) antibodies and incubated on ice in the dark for 1 h. After washing with PBS (30 s), the cells were tested on a BD FACSVerse flow cytometer. FlowJo v10 software was used for data analysis.

### T cell stimulation

The frozen cells were resuspended and washed with RPMI medium 1640 (Gibco), and then 1 × 10^6^ cells per well were transferred to a sterile 96-well plate and cultured with complete medium (RPMI 1640 supplemented with 10% FBS, penicillin, streptomycin). Then, 2 μg/ml RBD peptide pool (Sino Biological, PP002-A) was added to the medium for 18-h stimulation. Cells were stimulated with BSA as the negative control. Co-stimulators, i.e., 2 μg/ml anti-human CD28 (BD Biosciences, 1050151) and 100 KU/ml (sigma, I7908-10KU), were then added to the medium. In the last 6 h of stimulation, 5 μg/ml brefeldin A (BioLegend, 420601) and 2.5 μg/ml monensin (Selleckchem, S2324) were added to the medium. The cells were stimulated with leukocyte activation cocktail (BD Biosciences, 550583) as a positive control.

### Flow cytometry

The stimulated cells were washed with PBS containing 2% FBS. The cells were then stained on ice in the dark for 45 min with anti-CD3 (BD Biosciences, 557705), anti-CD4 (BioLegend, 317417), and anti-CD8 (BioLegend, 344712). The cells were fixed and permeabilized using a Cytofix/Cytoperm Kit (BD Biosciences, 554714). Afterwards, cells were stained with anti-IFN-γ (BioLegend, 502526), IL-2 (BioLegend, 500306), IL-4 (BioLegend, 500826), and IL-17A (BioLegend, 512329) antibodies and incubated on ice in the dark for 1 h. Cells were washed and resuspended with PBS and detected on the BD FACSVerse flow cytometer. FlowJo v10 was used for data analysis.

### Statistical analyses

All data are presented as means ± standard error of the mean (SEM), except for the titers of NAbs, which were quantified using geometric mean + geometric standard deviation. Student’s *t*-test and paired *t*-test were used to determine the statistical significance of differences between two groups. One-way analysis of variance (ANOVA) and two-way ANOVA were used to determine statistical significance for different dose groups and the curve graphs, respectively. Quantification graphs were analyzed using GraphPad Prism v8 (GraphPad Software). *: *P* < 0.05; **: *P* < 0.01; ***: *P* < 0.001.

## Acknowledgments

We thank all colleagues from the National Biosafety Laboratory (Wuhan), Chinese Academy of Sciences, China, for their support during the study. We thank the Center for Instrumental Analysis and Metrology and Biosafety Level 3 Laboratory, Wuhan Institute of Virology, Wuhan, China. We thank Professor Yifeng Zhou, Professor Li Bai, Dr. Fei Liu and their lab members for discussion and assisting in the design of the experiments. We thank Jia Wu, Jun Liu and Hao Tang from Wuhan Institute of Virology for the management of BSL-3 facility, where all the authentic SARS-CoV-2 experiments were conducted. This work was supported by Joint Laboratory of Innovation in Life Sciences from the University of Science and Technology of China (USTC) and Changchun Zhuoyi Biological Co. Ltd., Strategic Priority Research Program of the Chinese Academy of Sciences (XDA16020603 and XDB39000000), National Key Research and Development Program of China (2020YFA0112200), National Natural Science Foundation of China (81925009, 81790644, 81900855, 82000941), and Fundamental Research Funds for the Central Universities (WK5290000001, WK5290000002, WK2090050048, WK2070000174). The study was also supported by the Anhui Provincial Natural Science Foundation (1808085MH289 to M.Z.).

## Author contributions

T.X., Z.Y., T.J., and Z.Q. conceived the project and designed the experiments. D.T. and M.Z. developed the vaccines, tested their immunogenicity and safety, and wrote the manuscript. Y.Y. constructed the SARS-CoV-2 variants, coated the ELISA plates, and performed flow cytometry. H.X. and H.Z. performed the neutralizing antibody assay in Vero E6 cells. H.T., W.Z., M.L., Y.W., H.M., X.H., W.L., Y.C., Y.Y., Y.Y., K.L., S.S., Y.L., G.G., W.G., Y.P., S.C., J.R., J.Z., J.M., Q.Z., Y.Z., L.L., C.S., and K.Z. worked on data collection, analysis, and discussion. All authors edited and proofread the manuscript.

## Data and material availability

All data needed to evaluate the conclusions of the paper can be found in the manuscript and/or Supplementary Material. Additional data related to this paper may be requested from the lead contact Professor Tian Xue.

