## Supplemental File for "Single-dose AAV-based vaccine induces a high level of neutralizing antibodies against SARS-CoV-2 in rhesus macaques"

**Table S1. Summary of rhesus macaques for vaccination**

| Group | Animal NO. | Vaccination Dose | Gender | Age (Years) | Sacrificed time（dpv） | Experiment |
|---|---|---|---|---|---|---|
| 1 | 1H-1 | $1\times10^{12}$vg AAV-SRBD | M | 6 | Live | Elisa, competitive-ELISA, authentic virus neutralization, flow cytometry |
| 1 | 1H-2 | $1\times10^{12}$vg AAV-SRBD | M | 7 | Live | Elisa, competitive-ELISA, authentic virus neutralization, flow cytometry |
| 1 | 1H-3 | $1\times10^{12}$vg AAV-SRBD | F | 5 | Live | Elisa, competitive-ELISA, authentic virus neutralization, flow cytometry |
| 2 | 2H-1 | $1\times10^{12}$vg AAV-SRBD | F | 5 | 70 | Elisa, competitive-ELISA, body weight, pathological indicators in blood, hepatic function, flow cytometry |
| 2 | 2H-2 | $1\times10^{12}$vg AAV-SRBD | M | 6 | 70 | Elisa, competitive-ELISA, body weight, pathological indicators in blood, hepatic function, flow cytometry |
| 2 | 2H-3 | $1\times10^{12}$vg AAV-SRBD | M | 6 | 70 | Elisa, competitive-ELISA, body weight, pathological indicators in blood, hepatic function, flow cytometry |
| 2 | 2H-4 | $1\times10^{12}$vg AAV-SRBD | F | 5 | 70 | Elisa, competitive-ELISA, body weight, pathological indicators in blood, hepatic function, flow cytometry |
| 2 | 2M-1 | $1\times10^{11}$vg AAV-SRBD | M | 7 | 70 | Elisa, competitive-ELISA, body weight, pathological indicators in blood, hepatic function, flow cytometry |
| 2 | 2M-2 | $1\times10^{11}$vg AAV-SRBD | M | 5 | 70 | Elisa, competitive-ELISA, body weight, pathological indicators in blood, hepatic function, flow cytometry |
| 2 | 2M-3 | $1\times10^{11}$vg AAV-SRBD | F | 6 | 70 | Elisa, competitive-ELISA, body weight, pathological indicators in blood, hepatic function, flow cytometry |
| 2 | 2L-1 | $1\times10^{10}$vg AAV-SRBD | F | 6 | 70 | Elisa, competitive-ELISA, body weight, pathological indicators in blood, hepatic function, flow cytometry |
| 2 | 2L-2 | $1\times10^{10}$vg AAV-SRBD | M | 7 | 70 | Elisa, competitive-ELISA, body weight, pathological indicators in blood, hepatic function, flow cytometry |
| 2 | 2L-3 | $1\times10^{10}$vg AAV-SRBD | F | 5 | 70 | Elisa, competitive-ELISA, body weight, pathological indicators in blood, hepatic function, flow cytometry |
| 2 | 2C-1 | $1\times10^{12}$vg AAV-GFP | F | 5 | 70 | Elisa, competitive-ELISA, body weight, pathological indicators in blood, hepatic function, flow cytometry |
| 2 | 2C-2 | $1\times10^{12}$vg AAV-GFP | F | 6 | 70 | Elisa, competitive-ELISA, body weight, pathological indicators in blood, hepatic function, flow cytometry |
| 2 | 2C-3 | $1\times10^{12}$vg AAV-GFP | M | 6 | 70 | Elisa, competitive-ELISA, body weight, pathological indicators in blood, hepatic function, flow cytometry |

Figure. S1

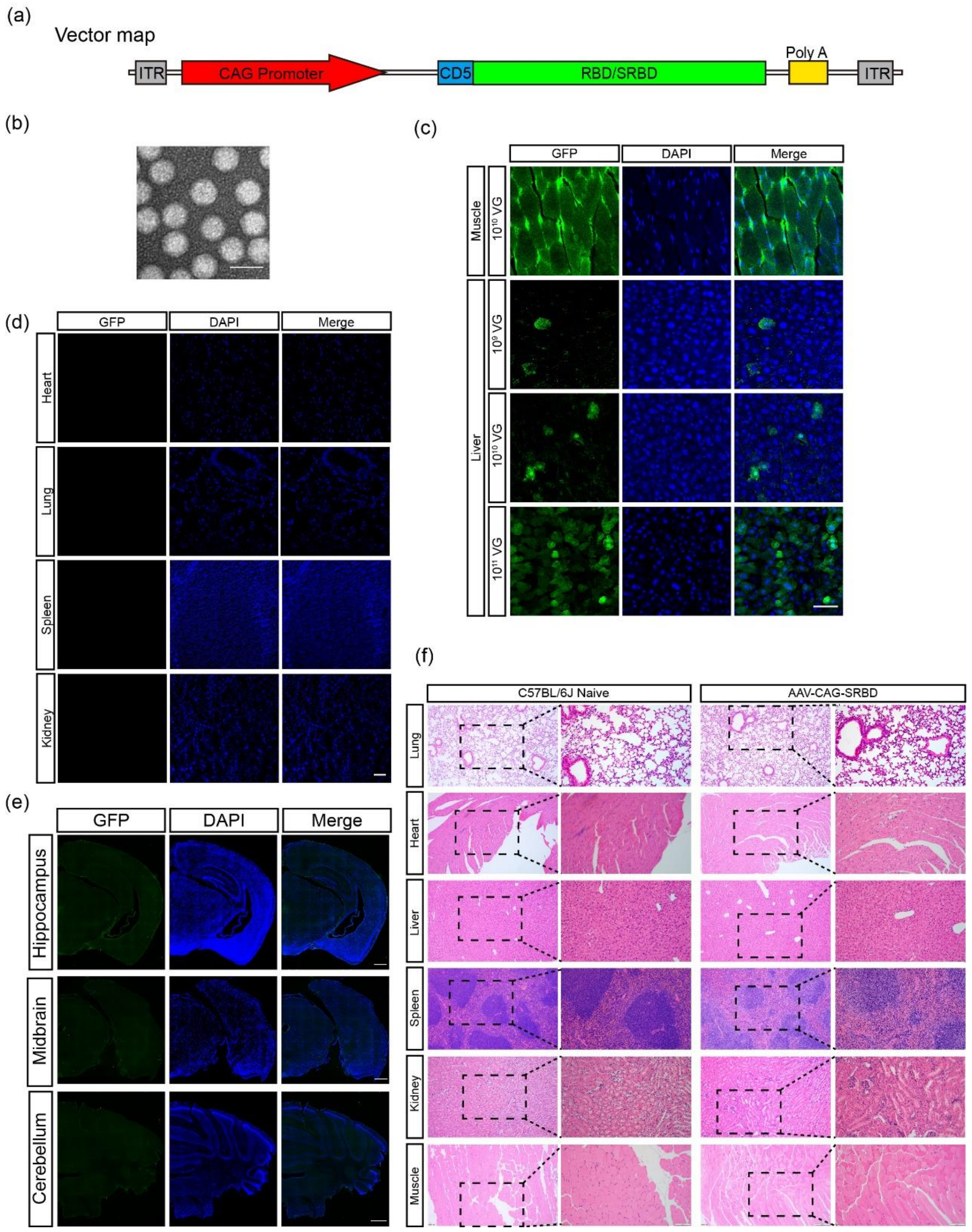


**Figure S1. Expression of AAV2/9- delivered GFP in the major organs of mice**

(a) Schematic representation of recombinant genome of AAV-SRBD and AAV-RBD vaccine candidates. CAG, CMV immediate enhancer/β-actin (CAG) promoter; ITR, inverted terminal repeat; CD5, CD5 signaling peptide.

(b) Images of AAV-SRBD particles by electron microscopy analysis (scale bar: 50 nm).

(c) Expression of GFP in muscle and liver from low/middle/high-dose AAV-GFP C57BL/6J mice at 42 dpv (scale bar: 50 μm).

(d) Expression of GFP in heart, lung, spleen, and kidney of high-dose AAV-GFP-injected C57BL/6J mice at 42 dpv (scale bar: 50 μm).

(e) Expression of GFP in brain of high-dose AAV-GFP-injected C57BL/6J mice at 28 dpv (scale bar: 500 μm).

(f) H&E-stained lung, heart, liver, spleen, kidney, and muscles of naïve (Control group) and AAV-SRBD-injected C57BL/6J mice at 42 dpv. Tissues were all normal in morphology (scale bar: 200 μm in the left row of each group and 100 μm in the right row of each group).

Figure. S2

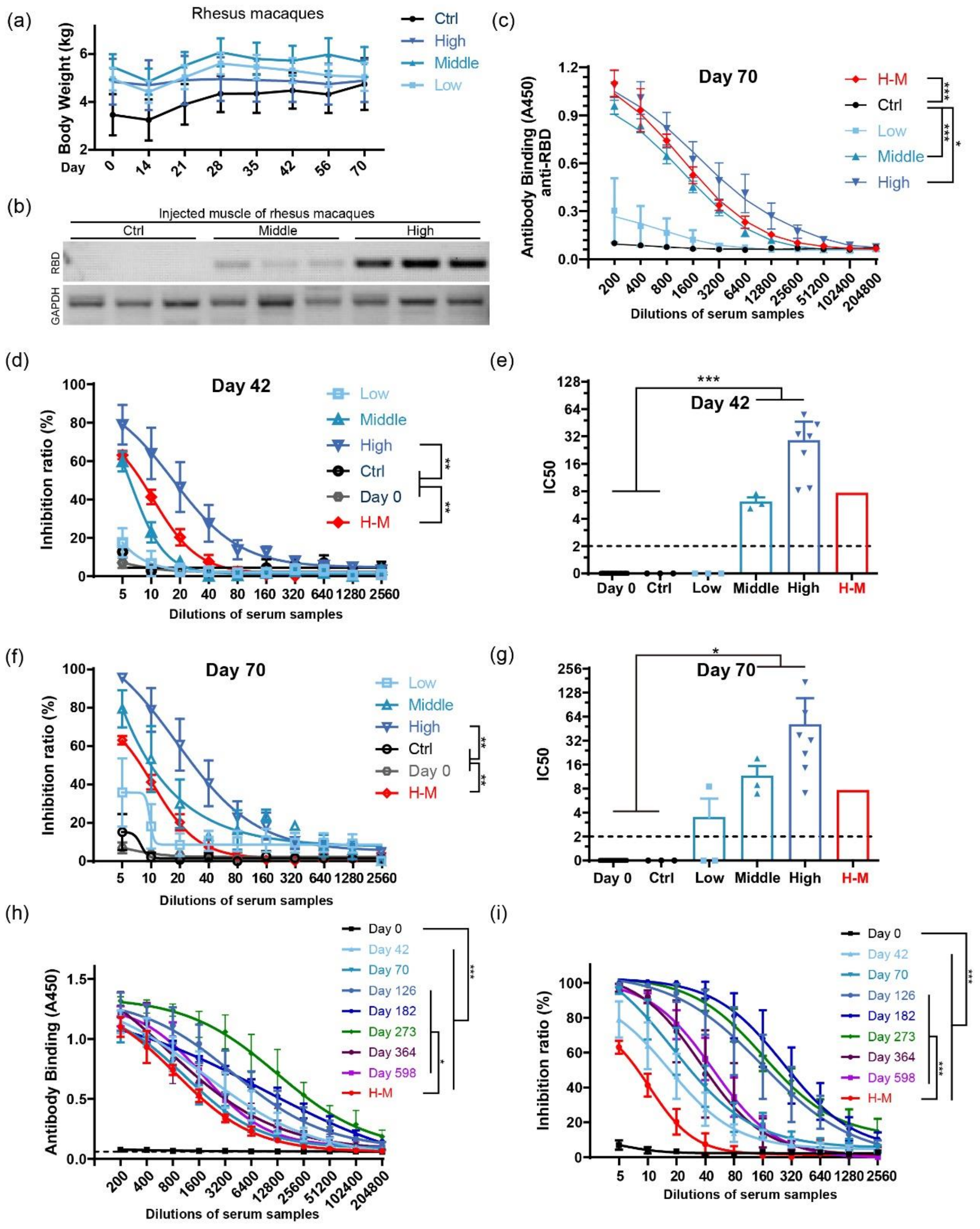


**Figure S2. Quantitative analysis of humoral responses in AAV-SRBD vaccine-injected rhesus macaques.**

(a) Weekly monitoring of body weight from 0 to 70 dpv in different dose macaques in group 2 (Ctrl: macaques intramuscularly injected with high-dose of AAV-CAG-GFP control; Low/Middle/High: macaques intramuscularly injected with low/middle/high-dose of AAV-SRBD vaccine; n = 3 macaques in Ctrl, Low, and Middle groups; n = 4 macaques in High group).

(b) RT-PCR data of RBD expression in middle- and high-dose AAV-SRBD- or AAV-CAG-GFP-injected macaques (n = 3 mice in each group).

(c) ELISA of RBD antibodies from macaque serum at 70 dpv (n = 3 macaques in Ctrl, Low, and Middle groups; n = 3 repeats in H-M group; n = 7 macaques in High group).

(d) Competitive ELISA of inhibition of SARS-CoV-2 RBD-hACE2 interaction by macaque serum at 42 dpv (n = 3 macaques in Ctrl, Low, and Middle groups; n = 3 repeats in H-M group; n = 7 macaques in High group; n = 16 macaques in Day 0 group).

(e) Quantitative analysis of RBD NAb half maximal inhibitory concentrations (IC50) between ACE2 and RBD, calculated by competitive ELISA at 42 dpv (n = 3 macaques in Ctrl, low, middle groups; n = 7 macaques in high-dose group; n = 16 macaques in Day 0 group). Mean IC50 of H-M is represented by red bar. The cutoff for the positive IC50 presented by black dot line.

(f) Competitive ELISA of inhibition of SARS-CoV-2 RBD-hACE2 interaction by macaque serum at 70 dpv (n = 3 macaques in Ctrl, Low, and Middle groups; n = 3 repeats in H-M group; n = 7 macaques in High group; n = 16 macaques in Day 0 group).

(g) Quantitative results of IC50 in e. Mean IC50 of H-M is represented by red bar. Positive cutoff of IC50 is represented by dotted line.

(h) ELISA of RBD antibodies from high-dose macaque serum in group 1 at 0, 42, 70, 126, 182, 273, 364 and 598 dpv (n = 3 macaques in each group; n = 3 repeats in H-M group).

(i) Competitive ELISA of inhibition of SARS-CoV-2 RBD-hACE2 interactions by high-dose macaque serum in group 1 at 0, 42, 70, 126, 182, 273, 364 and 598 dpv (n = 7 macaques in Day 0, 42, 70 group; n = 3 macaques in Day 126, 182, 273, 364, 598 group; n = 3 repeats in H-M group).

Values are means ± SEM or geometric mean + geometric standard deviation for antibody titer. *: $P < 0.05$; **: $P < 0.01$; ***: $P < 0.001$.

Figure. S3

PRNT50 of the high-dose macaque sera against
WT SARS-CoV-2 and variants

| Days \ Monkey NO. | | 1H-1 | 1H-2 | 1H-3 |
|---|---|---|---|---|
| Day 35 | WT | 60 | 1221 | 729 |
| | Beta | 250 | 3006 | 216 |
| | Delta | 70 | 2005 | 2344 |
| | Omi-cron | <20 | 89 | 57 |
| Day 56 | WT | 380 | 2498 | 1979 |
| | Beta | 450 | 4327 | 1036 |
| | Delta | 246 | 3483 | 3056 |
| | Omi-cron | 25 | 435 | 763 |
| Day 84 | WT | 1424 | 6803 | 8104 |
| | Beta | 2468 | 5394 | 3436 |
| | Delta | 611 | 20113 | 7112 |
| | Omi-cron | 104 | 2606 | 2528 |
| Day 98 | WT | 3808 | 6313 | 5362 |
| | Beta | 3839 | 13550 | 5176 |
| | Delta | 1229 | 13609 | 5371 |
| | Omi-cron | 249 | 4552 | 3194 |

| Days \ Monkey NO. | | 1H-1 | 1H-2 | 1H-3 |
|---|---|---|---|---|
| Day 182 | WT | 9891 | 7018 | 5342 |
| | Beta | 37161 | 14728 | 5590 |
| | Delta | 14837 | 5473 | 8137 |
| | Omi-cron | 10991 | 5423 | 3061 |
| Day 273 | WT | 16551 | 6313 | 7843 |
| | Beta | 45872 | 7133 | 1298 |
| | Delta | 7593 | 21988 | 9425 |
| | Omi-cron | 11112 | 5187 | 893 |
| Day 364 | WT | 5824 | >48600 | 4826 |
| | Beta | 32798 | >48600 | 2118 |
| | Delta | 7153 | >48600 | 13127 |
| | Omi-cron | 4625 | 5875 | 2450 |
| Day 598 | WT | 7348 | 5171 | 3751 |
| | Beta | 13319 | 5405 | 676 |
| | Delta | 4817 | 8921 | 5252 |
| | Omi-cron | 5974 | 4466 | 1132 |

**Figure S3. PRNT 50 values of high-dose macaques' sera in group 1 from 35 to 598 dpv against wild-type SARS-CoV-2, Beta, Delta and Omicron variants.**

Figure. S4

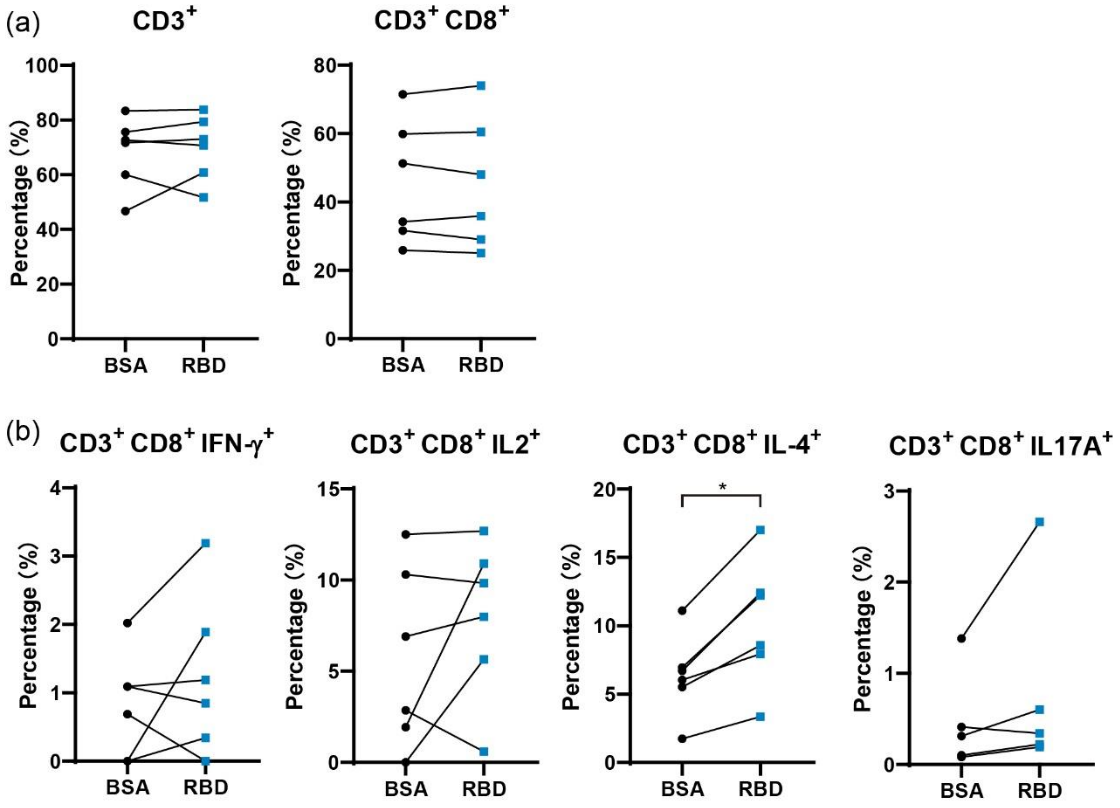


**Figure S4. Changes of immune cells after vaccination with AAV vaccine.**

(a) Percentage of $CD3^+$and $CD3^+$ $CD8^+$ cells in blood of high-dose rhesus macaques at day 35 dpv activated by BSA or RBD peptide.

(b) Percentage of $CD3^+$ $CD8^+$ IFN-$\gamma^+$, $CD3^+$ $CD8^+$ IL-$2^+$, $CD3^+$ $CD8^+$ IL-$4^+$ and $CD3^+$ $CD8^+$IL-$17A^+$ cells in blood of high-dose rhesus macaques at day 35 dpv activated by BSA or RBD peptide.

n = 6 macaques in each group, Values are means ± SEM. *: $P < 0.05$.

Figure. S5

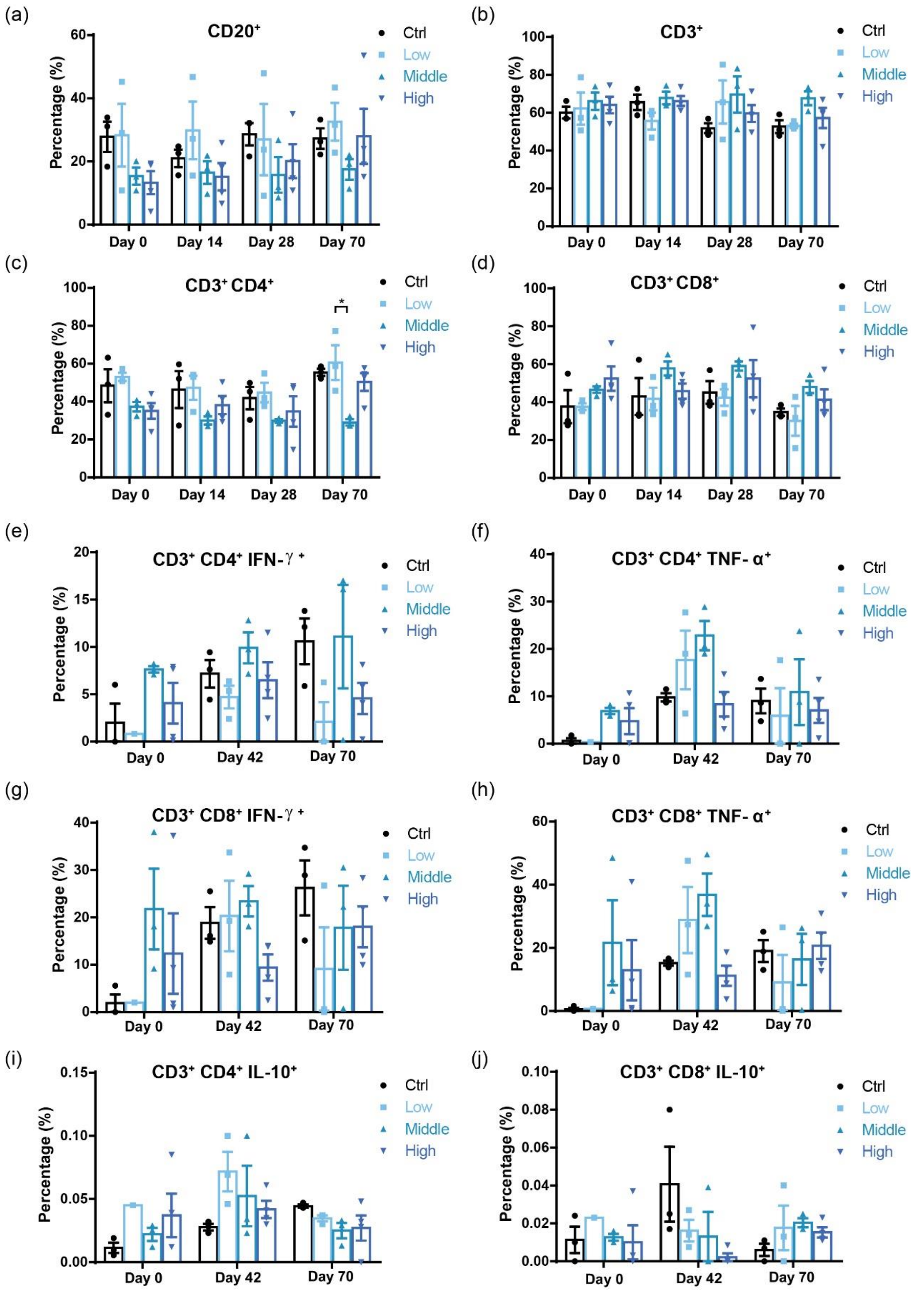


**Figure S5. Safety evaluation of SRBD vaccine with immune cells in rhesus macaques for 70 dpv.**

(a) Percentage of CD20$^+$ cells in serum of high/middle/low-dose and control rhesus macaques at 0, 14, 28 and 70 dpv (Ctrl: intramuscular injection with high-dose AAV-CAG-GFP control; Low/Middle/High: intramuscular injection of low/middle/high-dose SRBD vaccine).

(b) Percentage of $CD3^+$ cells in serum of high/middle/low-dose and control rhesus macaques at 0, 14, 28 and 70 dpv.

(c) Percentage of $CD3^+$ and $CD4^+$ cells in serum of high/middle/low-dose and control rhesus macaques at 0, 14, 28 and 70 dpv.

(d) Percentage of $CD3^+$ and $CD8^+$ cells in serum of high/middle/low-dose and control rhesus macaques at 0, 14, 28 and 70 dpv.

(e) Percentage of $CD3^+$, $CD4^+$, and IFN-$\gamma^+$ cells in serum of high/middle/low-dose and control rhesus macaques at 0, 42 and 70 dpv.

(f) Percentage of $CD3^+$, $CD4^+$, and TNF-$\alpha^+$ cells in serum of high/middle/low-dose and control rhesus macaques at 0, 42 and 70 dpv.

(g) Percentage of $CD3^+$, $CD8^+$, and IFN-$\gamma^+$ cells in serum of high/middle/low-dose and control rhesus macaques at 0, 42 and 70 dpv.

(h) Percentage of $CD3^+$, $CD8^+$, and TNF-$\alpha^+$ cells in serum of high/middle/low-dose and control rhesus macaques at 0, 42 and 70 dpv.

(i) Percentage of $CD3^+$, $CD4^+$, and IL-$10^+$ cells in serum of high/middle/low-dose and control rhesus macaques at 0, 42 and 70 dpv.

(j) Percentage of $CD3^+$, $CD8^+$, and IL-$10^+$ cells in serum of high/middle/low-dose and control rhesus macaques at s 0, 42 and 70 dpv.

n = 3 macaques in Ctrl, Low, and Middle groups; n = 4 macaques in High group, Values are means ± SEM. *: $P < 0.05$.

Figure. S6

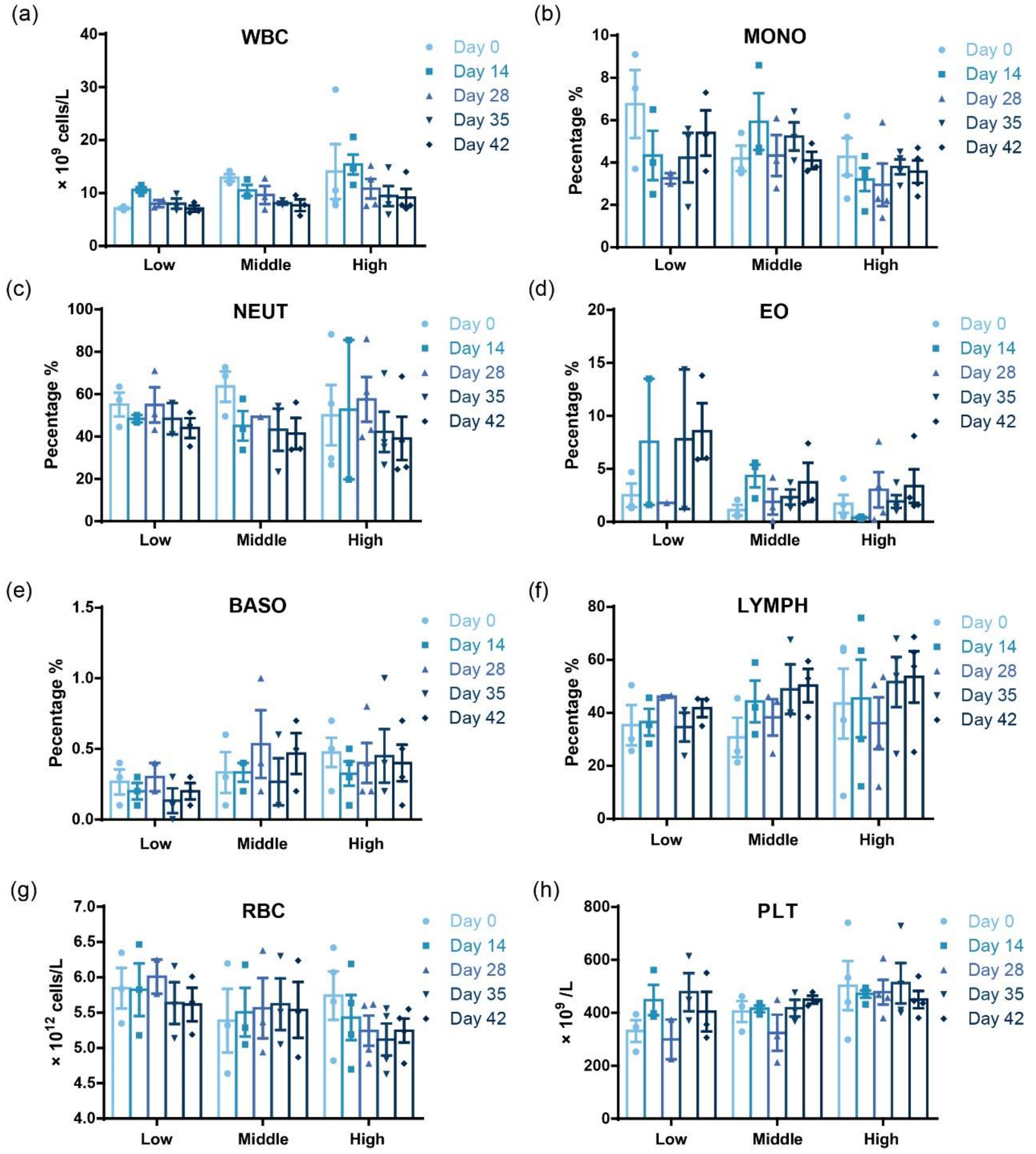


**Figure S6. Pathological indicators in blood of macaques following intramuscular injection of AAV-SRBD vaccine.**

(a) White blood cell (WBC) count in serum of high/middle/low-dose rhesus macaques at 0, 14, 28, 35, and 42 dpv.

(b) Monocyte cell (MONO) percentage in serum of high/middle/low-dose rhesus macaques at 0, 14, 28, 35, and 42 dpv.

(c) Neutrophil cell (NEUT) percentage in serum of high/middle/low-dose rhesus macaques at 0, 14, 28, 35, and 42 dpv.

(d) Eosinophil cell (EO) percentage in serum of high/middle/low-dose rhesus macaques at 0, 14, 28, 35, and 42 dpv.

(e) Basophil cell (BASO) percentage in serum of high/middle/low-dose rhesus macaques at 0, 14, 28, 35, and 42 dpv.

(f) Lymphocyte cell (LYMPH) percentage in serum of high/middle/low-dose rhesus macaques at 0, 14, 28, 35, and 42 dpv.

(g) Red blood cell (RBC) count in serum of high/middle/low-dose rhesus macaques at 0, 14, 28, 35, and 42 dpv.

(h) Platelet (PLT) count in serum of high/middle/low-dose rhesus macaques at 0, 14, 28, 35, and 42 dpv.

n = 3 macaques in Low and Middle groups; n = 4 macaques in High group, Values are means ± SEM

Figure. S7

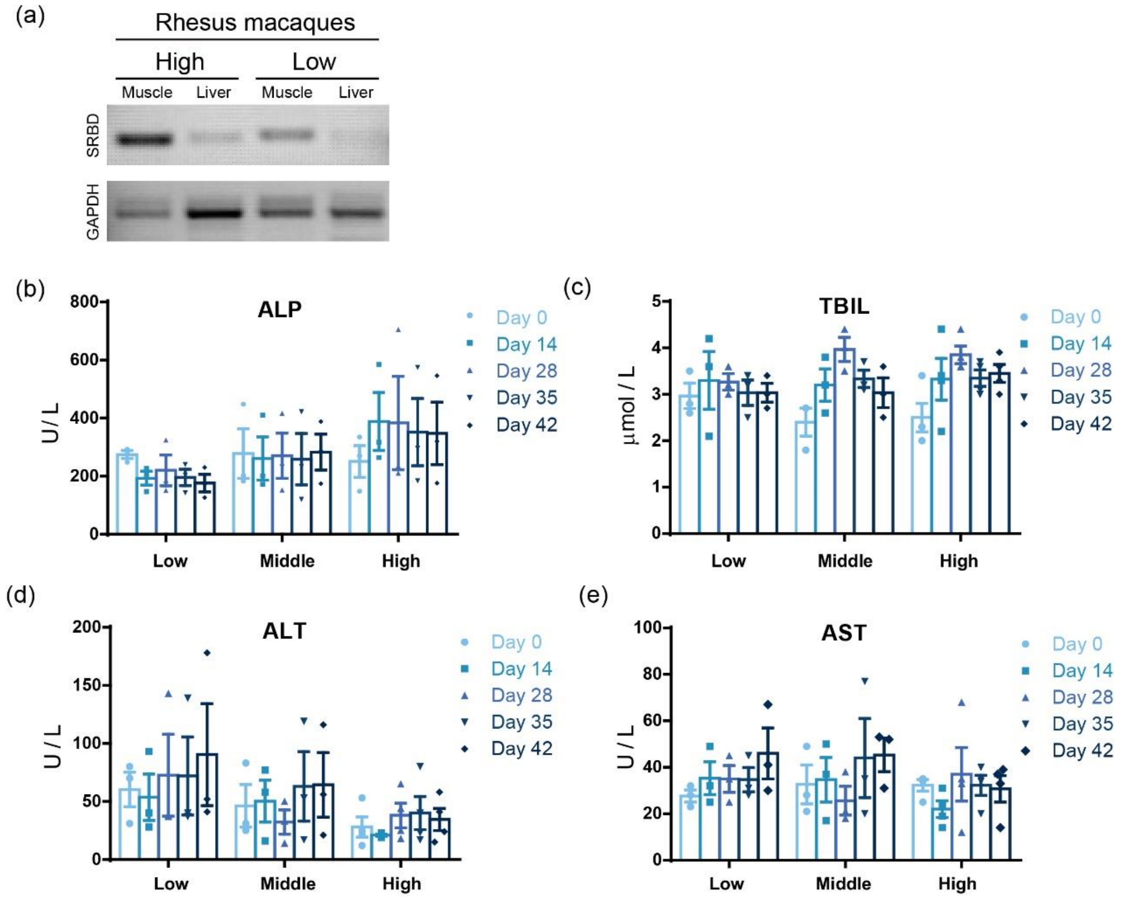


**Figure S7. Infection of AAV in muscles and liver of macaques and hepatic function of the immunized macaques.**

(a) RT-PCR of RBD in muscle and liver cells from high- and low-dose macaques (70 dpv).

(b) Alkaline phosphatase (ALP) level in serum of high/middle/low-dose rhesus macaques at 0, 14, 28, 35, and 42 dpv.

(c) Total bilirubin (TBIL) level in serum of high/middle/low-dose rhesus macaques at 0, 14, 28, 35, and 42 dpv.

(d) Alanine aminotransferase (ALT) level in serum of high/middle/low-dose rhesus macaques at 0, 14, 28, 35, and 42 dpv.

(e) Aspartate aminotransferase (AST) level in serum of high/middle/low-dose rhesus macaques at 0, 14, 28, 35, and 42 dpv.

n = 3 macaques in Low and Middle groups; n = 4 macaques in High group, Values are means ± SEM

Figure. S8

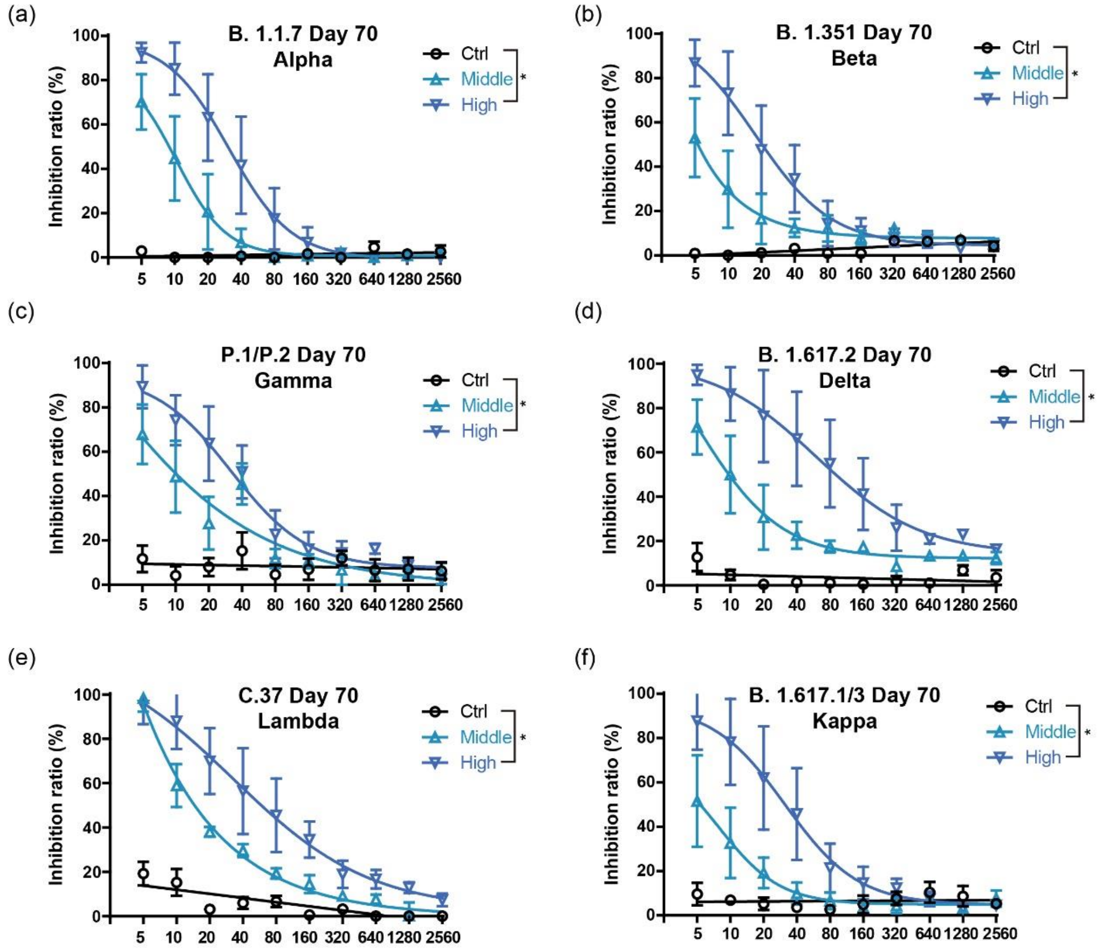


**Figure S8. AAV-SRBD immune serum efficiently inhibited the binding of ACE2 with RBD variants.**

(a) Competitive ELISA of inhibition of RBD domain in Alpha variant and hACE2 interactions by macaque serum at 70 dpv. (Middle/High: 70 dpv of middle/high-dose SRBD vaccine; Ctrl: 70 dpv of high-dose AAV-CAG-GFP control; n = 3 macaques in each group).

(b) Competitive ELISA of inhibition of RBD domain in Beta variant and hACE2 interactions by macaque serum at 70 dpv (n = 3 macaques in each group).

(c) Competitive ELISA of inhibition of RBD domain in Gamma variant and hACE2 interactions by macaque serum at 70 dpv (n = 3 macaques in each group).

(d) Competitive ELISA of inhibition of RBD domain in Delta variant and hACE2 interactions by macaque serum at 70 dpv (n = 3 macaques in each group).

(e) Competitive ELISA of inhibition of RBD domain in Lambda and hACE2 interactions by macaque serum at 70 dpv (n = 3 macaques in each group).

(f) Competitive ELISA of inhibition of RBD domain in Kappa variant and hACE2 interactions by macaque serum at 70 dpv (n = 3 macaques in each group).

Values are means ± SEM. *: $P < 0.05$.
